# Beyond a ribosomal RNA methyltransferase, the wider role of MraW in DNA methylation, motility and colonization in *Escherichia coli* O157:H7

**DOI:** 10.1101/480244

**Authors:** Xuefang Xu, Heng Zhang, Ying Huang, Yuan Zhang, Xiaoyuan Wang, Dai Wang, Ji Pu, Hongqing Zhao, Xuancheng Lu, Shuangshuang Lu, Yanwen Xiong, Changyun Ye, Yuhui Dong, Ruiting Lan, Jianguo Xu

## Abstract

MraW (RsmH) is an AdoMet-dependent 16S rRNA methyltransferase conserved in bacteria and plays a role in the fine-tuning of the ribosomal decoding center. It was recently found to contribute to the virulence of *Staphylococcus aureus* in host animals. In this study, we examined the function of MraW in *Escherichia coli* O157:H7 and found that deletion of *mraW* led to decreased motility and flagellar production. Whole-genome bisulfite sequencing showed genome wide decrease of methylation of 336 genes and 219 promoters in the *mraW* mutant. The methylation level of 4 flagellar gene sequences were further confirmed by bisulfite PCR sequencing. Quantitative reverse transcription PCR results indicated the transcription of these genes was also affected. MraW was observed to directly bind to the four flagellar gene sequences by electrophoretic mobility shift assay (EMSA). A common motif in differentially methylated regions of promoters and coding regions of the 4 flagellar genes was identified. Reduced methylation was correlated with altered expression of 21 of the 24 genes tested. DNA methylation activity of MraW was confirmed by DNA methyltransferase (DNMT) activity assay *in vitro*. The *mraW* mutant colonized poorer than wild type in mice. we further found that the expression of *mraZ* in the *mraW* mutant was increased confirming the antagonistic effect of *mraW* on *mraZ*. In conclusion, *mraW* was found to be a DNA methylase and has a wide-ranging effect on *E*. *coli* O157:H7 including motility and virulence *in vivo* via genome wide methylation and *mraZ* antagonism.

**IMPORTANCE:** MraW is a well-studied 16S rRNA methyltransferase and was recently found have an impact on bacterial virulence. Here we demonstrated its new function as a DNA methylase and effect on motility, colonization in mice, DNA methylation in genome wide and contribution to virulence. Its direct binding of differentially methylated flagellar-encoding DNA sequences was observed, indicating a correlation between DNA methylation and regulation of flagellar genes. In addition, the expression of *mraZ* which function as an antagonist of *mraW* was increased in the *mraW* mutant. *mraW* plays an important role in gene regulation likely through DNA methylation. Clearly it plays a role in virulence in *E. coli* O157:H7. It also opens a new research field for virulence study in bacteria.

---

*Escherichia coli* O157:H7 is the most commonly isolated enterohaemorrhagic *E*. *coli* (EHEC) and accounts for more than 90% of clinical EHEC cases (1–3). It causes diarrheal diseases and other syndromes such as hemorrhagic colitis (HC) and hemolytic uremic syndrome (HUS) with colonization of the intestinal mucosa and subsequent toxin release in the intestinal tract (4). The main virulence factors involved in intestinal colonization of the host are the type III secretion system (T3SS), curli and flagella (5, 6). The regulations of the T3SS and flagella are very complex and affected by many regulators and environmental factors such as pH value, glucose, iron and temperature (3, 7–9). Recently, it was reported that the utilization of carbon nutrition can affect colonization of *E. coli* in the mouse intestine (10, 11).

MraW (or named as RsmH) is a 16S RNA methyltransferase (MTase) responsible for N4-methylation of C1402 in bacteria, which is also methylated by another MTase YraL (or named as RsmI) 2’-O-methylation (m^4^Cm) (12). Recent studies have shown that *mraW* participates in virulence. It was revealed that both *rsmI* and *rsmH* (*mraW*) affected the virulence of *Staphylococcus aureus* in silk worms by contributing resistance to oxidative stress (13).

DNA methylation has been well studied in eukaryote and is essential in the development and progression of cancer (14). It becomes a rapidly growing area of research due to its contribution to improved diagnosis and treatment. However, methylation effect on virulence in bacteria has not been well studied and remains unknown. We hypothesized that methylation in bacteria affect the virulence as it does in eukaryotes. The aim of this study was to determine the effect of *mraW* on virulence in enterohaemorrhagic *E. coli* O157:H7 and the relationship between methylation and virulence.

## RESULTS

### Deletion of *mraW* led to reduced motility and flagellin production/secretion

The gene encoding MraW was deleted in *E. coli* O157:H7 strain EDL933 resulting in a mutant strain designated as EDL933ΔmraW. The motility of the wild type and the mutant was determined by the radius of chemotactic ring which was 2.3 ± 0.1 cm and 0.783 ± 0.03cm respectively. The difference is statistically significant (t test, P < 0.01) (Fig. 1. A-B). The decrease in motility in the *mraW* mutant was also correlated with a decrease in the production or secretion of FliC as determined by western blotting using anti-H7 antisera (Fig. 1C). To determine whether the *mraW* deletion can be complemented, we created pBADmraW, a low copy number plasmid carrying *mraW* which was transformed into EDL933ΔmraW. The radius of the chemotactic ring of the complemented strain (EDL933ΔmraW+pBADmraW) was 2.55 ± 0.09 cm (Fig. 1A), which is similar to the wild type. Thus, the decrease in motility of the mutant was almost complemented back by the plasmid pBADmraW expressing *mraW*. The generation time of EDL933, EDL933ΔmraW and EDL933ΔmraW+pBADmraW was 35.7 ± 0.06 min, 35.9 ± 0.04 min and 35.8 ± 0.1 min respectively. Hence the difference in motility and production of FliC was not due to the growth rate which was similar among the mutant, the complemented strain and the wild type.

**Figure 1.**
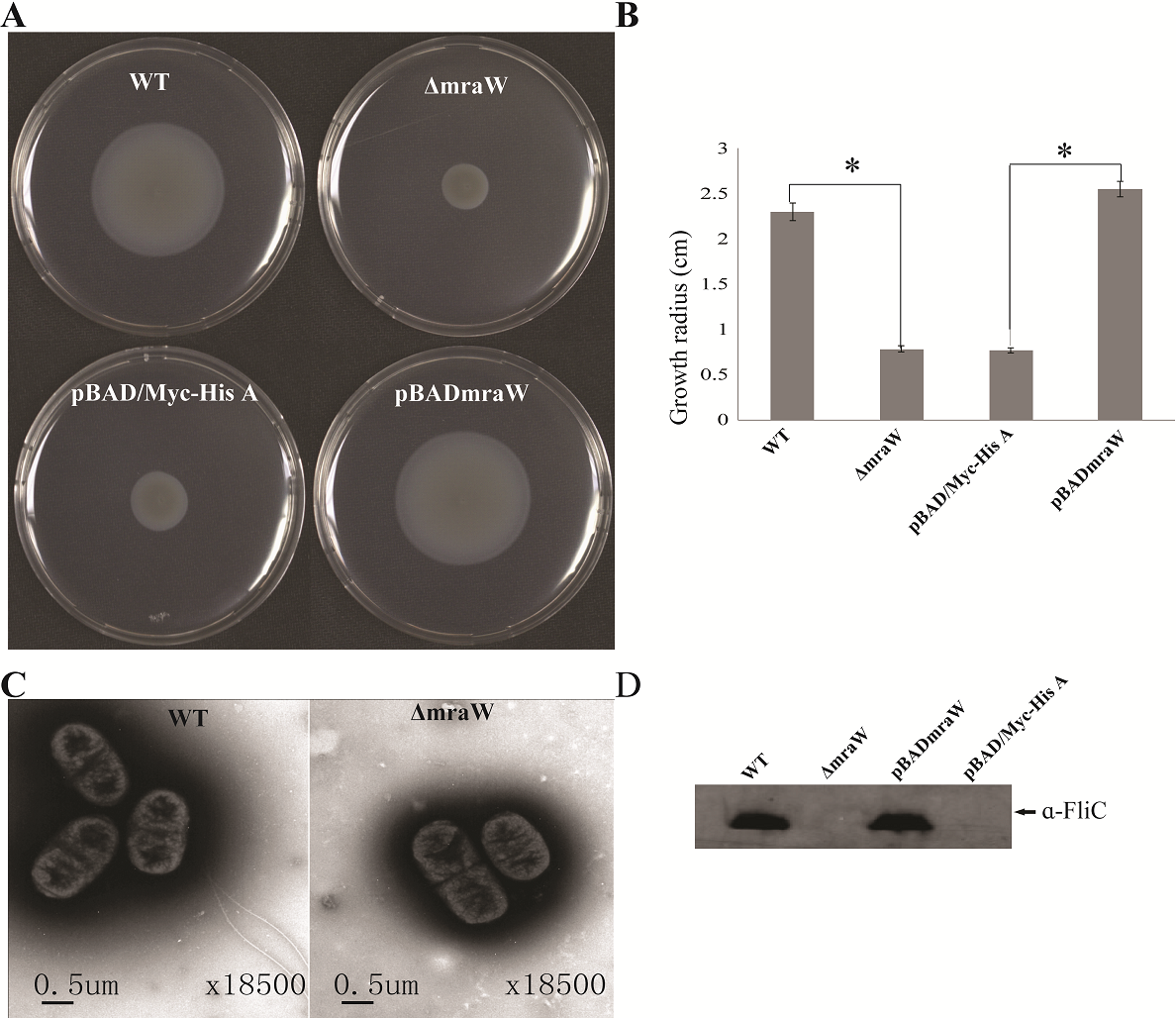
Effects of *mraW* on motility. (A, B) Representative images and growth radius of swimming motility for the wild-type EDL933, the *mraW* deletion mutant (EDL933ΔmraW), and complemented strain (EDL933ΔmraW +pBADmraW). pBAD/Myc-His A is an empty vector control. Error bar shows the standard deviation from three independent experiments. Differences were analyzed for significance by using T-test. Significant difference between two strains (P <0.01) isindicated by a * with a linked line. (C) Transmission electron micrographs of wild-type EDL933 and the *mraW* mutant (scale bar, 0.5 μm). (D) Immunoblot analysis of FliC protein in the whole cell lysates prepared from wild-type EDL933, the *mraW* deletion mutant (EDL933ΔmraW), empty vector control strain (EDL933ΔmraW+pBAD/Myc-His A) and complemented strain (EDL933ΔmraW+pBADmraW) grown in LB. Arrows indicate a reactive band corresponding to FliC detected with anti-H7 FliC antibodies. Figure 2.

To investigate whether the reduced motility of the EDL933mraW mutant was due to decrease of surface flagella, bacteria were inspected by transmission electron microscopy (TEM) to visualize surface flagella at a magnification of 9,700X. Thirty fields of view were randomly selected and about 50-200 cells counted. The majority (95%) of the *mraW* deletion mutants were found to have no flagella, with the remaining 5% of the cells having 1 to 2 flagella (Fig. 1D). In contrast, most of the cells from the wild-type EDL933 had surface flagella although the number of surface flagella was limited to between one and three. The electron microscope results suggest that the expression of flagella has diminished in the *mraW* deletion mutant under the culture condition used in this study.

### The effect of *mraW* on genome wide DNA methylation

Besides the function as a methyltransferase targeting16S RNA, we investigated whether MraW affects DNA methylation at the genome level. Genome-wide methylation profiling was performed in wild-type EDL933 and EDL933ΔmraW by bisulfite sequencing to obtain detailed information on the methylation status of each cytosine. A total of 1.2 Gigabytes of sequence data were obtained for each strain. The sequencing depth of EDL933 and EDL933Δmra was 104.24 fold and 99.96 fold. The methylation level of C, CG, CHG, CHH (where H = A, T or C) in EDL933 and EDL933ΔmraW whole genomes was 1.56, 1.09, 3.84 and 0.44 and 1.42, 0.96, 3.50 and 0.41 respectively (see Table S1 in the supplemental material). Although a similar methylation level was found in cytosine in EDL933ΔmraW compared to EDL933, there was a trend of differences in methylation levels in genes and promoter regions (500 bp upstream of the coding regions) which contained at least one differentially methylated regions (DMRs). The methylation levels of 219 promoters including 97 with known function and 336 genes including 152 with known function (FDR < 0.05,p<0.005) (Fig. 2A-B) in EDL933ΔmraW were lower compared with the wild type. These differentially methylated promoter regions and genes are discussed in detail below.

**Figure 2.**
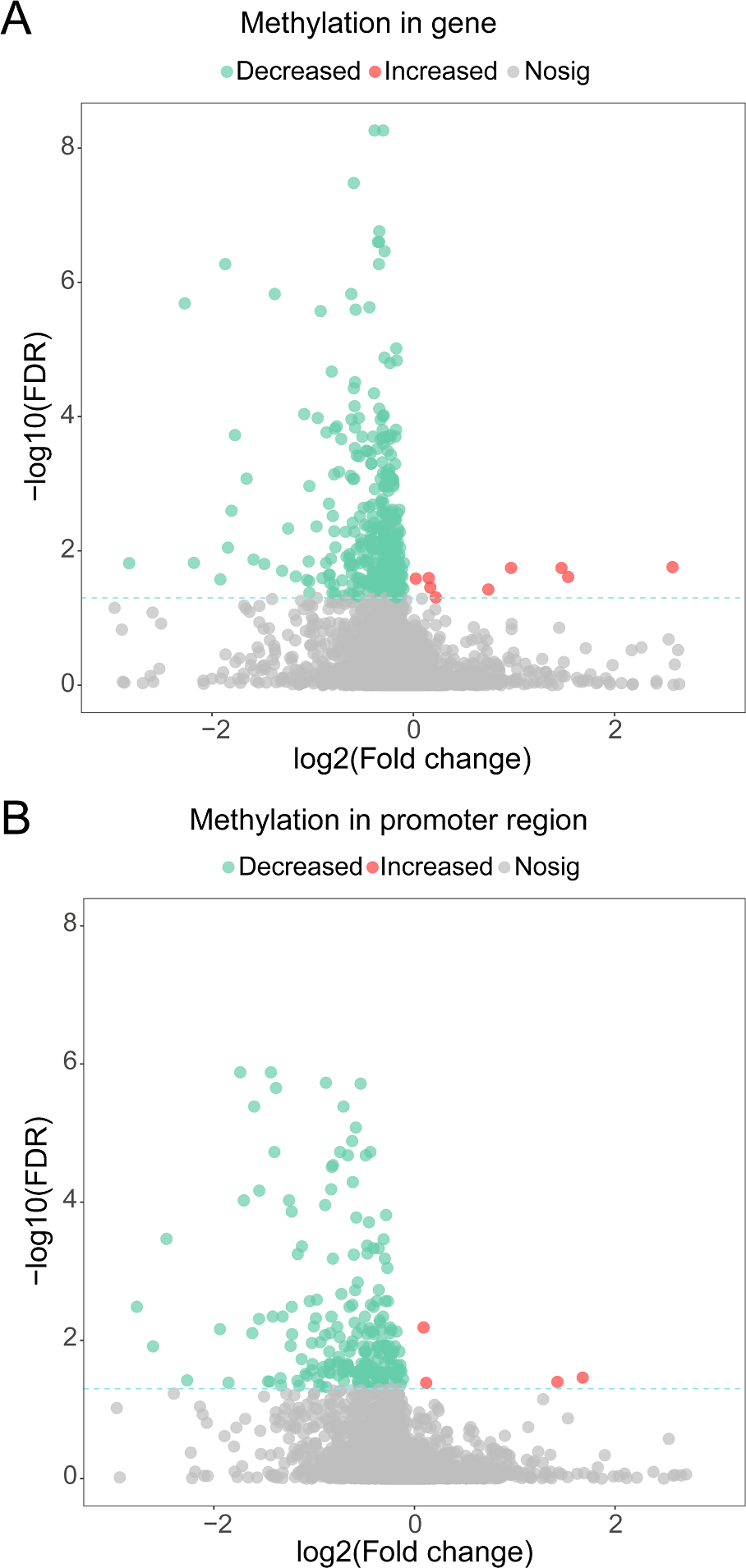
Differential methylation between EDL933 and EDL933ΔmraW. (A) The methylation difference between EDL933 and EDL933ΔmraW in genes. The log2 difference (EDL933ΔmraW versus EDL933) of methylation (x-axis) by −log10 (false discovery rate [FDR]) (y-axis) is shown. The broken green line shows the significance cutoff corresponding to an FDR less than 0.05. Red spots indicate genes with increased methylation; green spots indicate genes with decreased methylation. (B) The methylation difference between EDL933 and EDL933ΔmraW in promoter regions. The log2 difference (EDL933ΔmraW versus EDL933) of methylation (x-axis) by −log10 (FDR) (y-axis) is shown. The broken green line shows the significance cutoff corresponding to an FDR less than 0.05. Red spots indicate promoter regions with increased methylation; green spots indicate promoter regions with decreased methylation.

### The effect of *mraW* on the methylation of flagella, energy metabolic pathway and other virulence genes

Consistently with decreased motility and flagellin secretion in the *mraW* mutant, two flagellar genes, *fliJ* (encoding a cytoplasmic flagellar protein) and *fliR* (flagellar apparatus integral membrane protein) and two promoters of flagellar genes, *fliK* (flagellar hook length) and *fhiA* (flagellar apparatus integral membrane protein) showed a lower level of methylation in the mutant compared to the wild type (Table S2 and S3).

In addition, the methylation level of the promoter of *qseB*, a flagellar related quorum sensing regulator gene, was decreased in the *mraW* mutant. QseBC regulates flagella indirectly as an enhancer of flagellar master regulator FlhDC and in the absence of QseC, phosphorylation of QseB can act as a repressor of the flagellar expression (15). In addition to flagella related genes, the largest proportion of DMRs are related to energy metabolism pathways. 30% (29/97) promoters and 26% (39/152) genes with DMRs participate in energy metabolism accounting for the highest proportion in the whole genome DMRs (Table S2 and S3). The methylation level of *barA* was also reduced which encodes one of the members of the BarA-UvrY two component system and plays an essential function in metabolic adaptation by controlling the carbon storage regulation system, Csr (16).

Many virulence genes and promoters in the *mraW* mutant were also affected at methylation level (Table S2 and S3), including three genes of the T3SS (*escD, espB* and *z4187*), three genes encoding three heat shock proteins (HtrC, Ddg and GrpE), a clod shock protein gene (*cspA*), a helicase gene (*recD*), and a curli activator gene (*crl*). A very small number of genes/promoters (14) had increased methylation level in the *mraW* mutant (Fig. 2A-B). Most of these genes were of unknown function. The five genes with known functions were *z1799*-encoding a prophage CP-933N encoded membrane protein, *tyrP*-tyrosine-specific transport system, *cspA*-transcriptional activator of HNS, *frwD*-PTS system fructose-like IIB component 2 and *eno*-enolase.

### Validation of bisulfite sequencing results

To validate the genome bisulfate sequencing results, 9 regions(4 genes and 5 promoters)including *fliR, fliJ, z2975, PfliK, PfhiA, PyidP, PtreR*, *z1440* and *z4981* were selected for BSP. Among these DMRs, *z1440* and *z4981* showed increased methylation level. The methylation levels were higher by BSP in both the wild type and the *mraW* mutant, but the magnitude of reduction of methylation (38% reduction on average) is similar to genome bisulfate sequencing results (30% reduction on average). These BSP data analyses revealed highly similar methylation patterns compared with genome bisulfate sequencing results including DMRs with increased methylation level (Table S4).

### Motifs of DMRs and MraW binding to DMRs of flagellar gene promoters and coding sequences

Given the methylation effect is genome wide, we searched for common motifs in the DMRs of the whole genome. No common motif with a p value < 0.0001 was found. We then restricted the searches to the DMRs of the flagellar genes and heat shock protein genes. In the four flagella genes, three genes *fliJ, fliK, fhiA* containing 6 DMRs while *fliR* had no DMR. One motif, GATGAAAGGC, was common in these 6 DMRs with a p value < 0.0001 (Fig. 3A). Similarly, common motifs in 21 DMRs of the three heat shock protein genes *htrC, ddg* and *grpE* were investigated. One common motif, ATTACT, was found with p value < 0.0001 (Fig. 3A). Since ATTACT occurs 1330 times in the EDL933 genome, we did not pursue this motif further.

**Figure 3.**
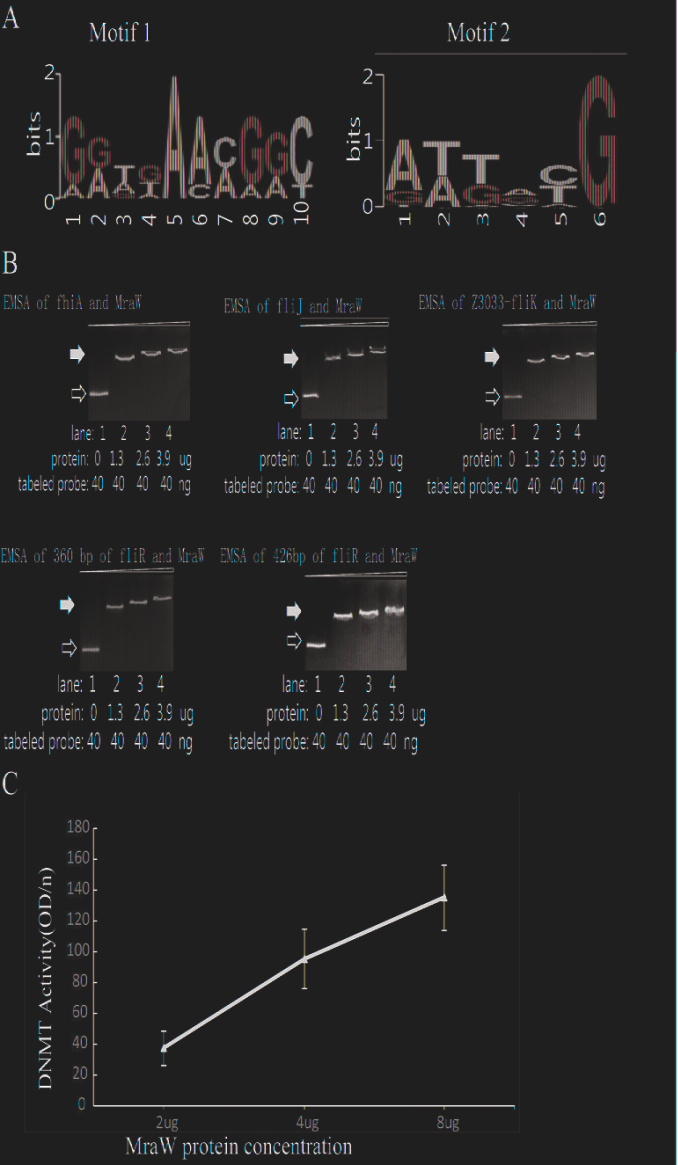
The methylation effect of MraW and binding interaction with flagellar genes or promoter regions. (A) DNA binding motif of *mraW*. DNA binding motif of mraW in flagellar gene related DMRs was analyzed using MEME program and a common motif from 6 DMRs was found: GGTGAACGGC (left). DNA binding motif of *mraW* in heat shock protein gene DMRs was analyzed using MEME program and a common motif from 21 DMRs was found: ATTACG (right). (B) Assessment of MraW binding to the DMRs of flagellar gene and promoter fragments. Reaction constituents are indicated underneath. Open arrow indicates free labelled DNA, the black arrow indicates MraW-DNA complexes. (C) MraW DNA methylation activity. DNMT activity increased with higher concentration of MraW proteins.

The DMRs of flagellar sequences and potential motifs clearly indicated the possibility of physical interaction between MraW and flagellar sequences. Theerfore, MraW was expressed and purified for electrophoretic mobility shift assay to assess its binding ability. Four flagellar nucleotide sequences namely, promoters of *fhiA* & *fliK*, and coding sequences of *fliJ* & *fliR* including the motif were cloned and used for the assay (Table S5). Three protein concentrations were tested. Binding occurred at the lowest concentration with major shift of mobility. Increasing concentrations of MraW led to small decreases in mobility (Fig. 3B). The binding results suggest that MraW could directly affect the expression of these flagellar genes.

### DNA methyltransferase (DNMT) activity of MraW

Since MraW can directly bind to the flagellar gene sequences, we tested whether MraW methylates DNA. A commercial ELISA kit containing DNMT substrate (cytosine) and anti-5-methylcytosine antibody was used for DNMT activity measurement. MraW protein with different concentrations was added and incubated with the DNMT substrate. Methylated DNA can be recognized with the anti-5-methylcytosine antibody which was attached with an enzyme catalyzing the substrate to blue color. DNMT activity of MraW ranged from 37 OD/h to 137 OD/h when the MraW protein concentration increased from 2 µg to 8 µg (Fig. 3C). Hence there is a positive correlation between the DNMT activity and MraW protein concentration indicating a DNA methylation function of MraW.

### Influence of *mraW* on colonization in mice

We next asked whether *mraW* would affect the colonization of EDL933 *in vivo* using the mouse model. To assess the virulence effect *in vivo*, a constitutively luminescent plasmid was introduced into the wild type and the mutant to identify them in the mixed infection experiments. Six-week-old female BALB/c mice were used. Two groups of 10 mice were intragastrically administered approximately 10^9^ and 10^10^ CFU of equal mix of wild type EDL933 and the *mraW* deletion mutant. We used two different inoculum sizes as better colonization was observed in high inoculum (above 10^9^ CFU) in previous studies (17, 18). The maintenance of the bioluminescent plasmid was assessed by luciferase scanning. The degree of bacterial colonization was measured by the number of luminescent bacteria present per gram feces. Mice was fed the same numbers of bacteria of the EDL933 and EDL933ΔmraW. Fecal shedding was examined up to day 7. Each colony from the shedding fecal sample was luminescent on Sorbitol–MacConkey agar plates with ampicillin for selection of *E. coli* O157:H7 (Fig. 4A). The *mraW* deletion mutant was selected by kanamycin resistance. A very similar shedding level of these two strains were found at day 1. A significant difference in colonization between EDL933 and EDL933ΔmraW was found from day 2 to day 7 (Fig. 4B-C). A 100 times difference was found between the wild type and the *mraW* deletion mutant from day 2 in both 10^9^ and 10^10^ CFU inoculum groups. The results indicated that the *mraW* deletion mutant colonized the mice poorer than the wild type EDL933.

**Figure 4.**
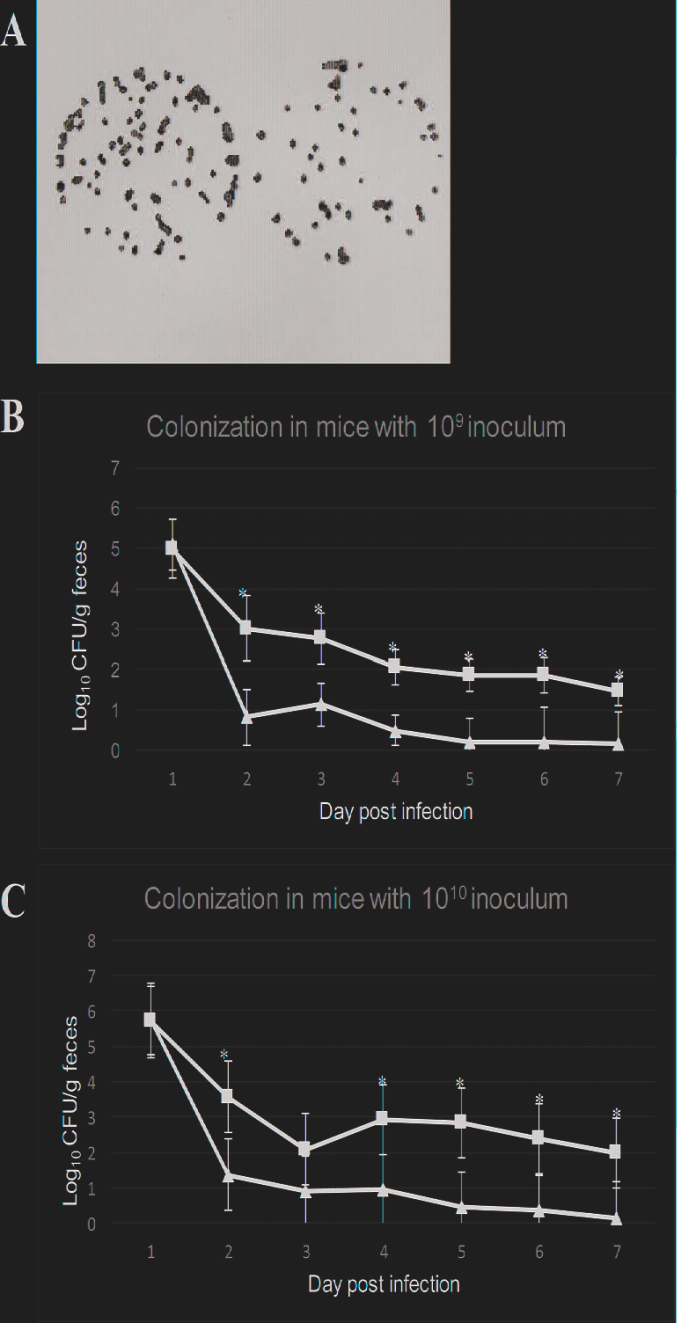
Colonization of BALB/c mice by EDL933 and EDL933ΔmraW with luminescent plasmid pGEN-luxCDABE. (A) Results showed that each colony contained luminescent plasmid pGEN-luxCDABE. (B) Colonization levels of EDL933 and EDL933ΔmraW over 7 days after oral gavage with a 10^9^ CFU mixed inoculum (dashed lines with ▪ EDL933 and ▴ for the *mraW* mutant. (C) Colonization levels of EDL933 and EDL933ΔmraW over 7 days after oral gavage with a 10^10^ CFU mixed inoculum (dashed lines with ▪ EDL933 and ▴ for the *mraW* mutant).

### The effect of *mraW* on the expression of a selected set of genes

We investigated the effect of *mraW* deletion on a selected set of genes which are related to virulence phenotype, primarily based on methylation effect detected above. The mRNA expression of flagella related genes *fliJ, fliR, fliK* and *fhiA* were all decreased (t test, P < 0.01) (Fig. 5A), which is consistent with the decline of flagellin protein production/secretion. Curli related gene, *crl* and the helicase gene, *recD*, were also decreased at the transcriptional level (t test, P < 0.01) (Fig. 5A). However, three genes encoding heat shock proteins, *htrC, ddg* and *grpE*, were significantly increased at the transcriptional level (t test, P < 0.01) (Fig. 6B). The mRNA transcription of cold shock protein *cspA* was also increased in a low but significant level (t test, P < 0.01) (Fig. 5C). Nine genes of carbon nutrition metabolism relevant to colonization were examined with qRT-PCR (10, 11). As expected, *rbsD, rbsA, rbsK, nanA, gmD, fcI, agaI-2, galP, fucO, araJ*, and *kdgK* all showed a reduced mRNA expression (t test, P < 0.01) (Fig. 5C). Type III secretion phenotype was investigated in the mutant. T3SS proteins were compared between the wild type and the mutant. No significant difference was found in protein level between these two strains (Fig.5D). Although the transcription levels of *escD, espB* and *z4187* were all increased (Fig. 5D).

**Figure 5.**
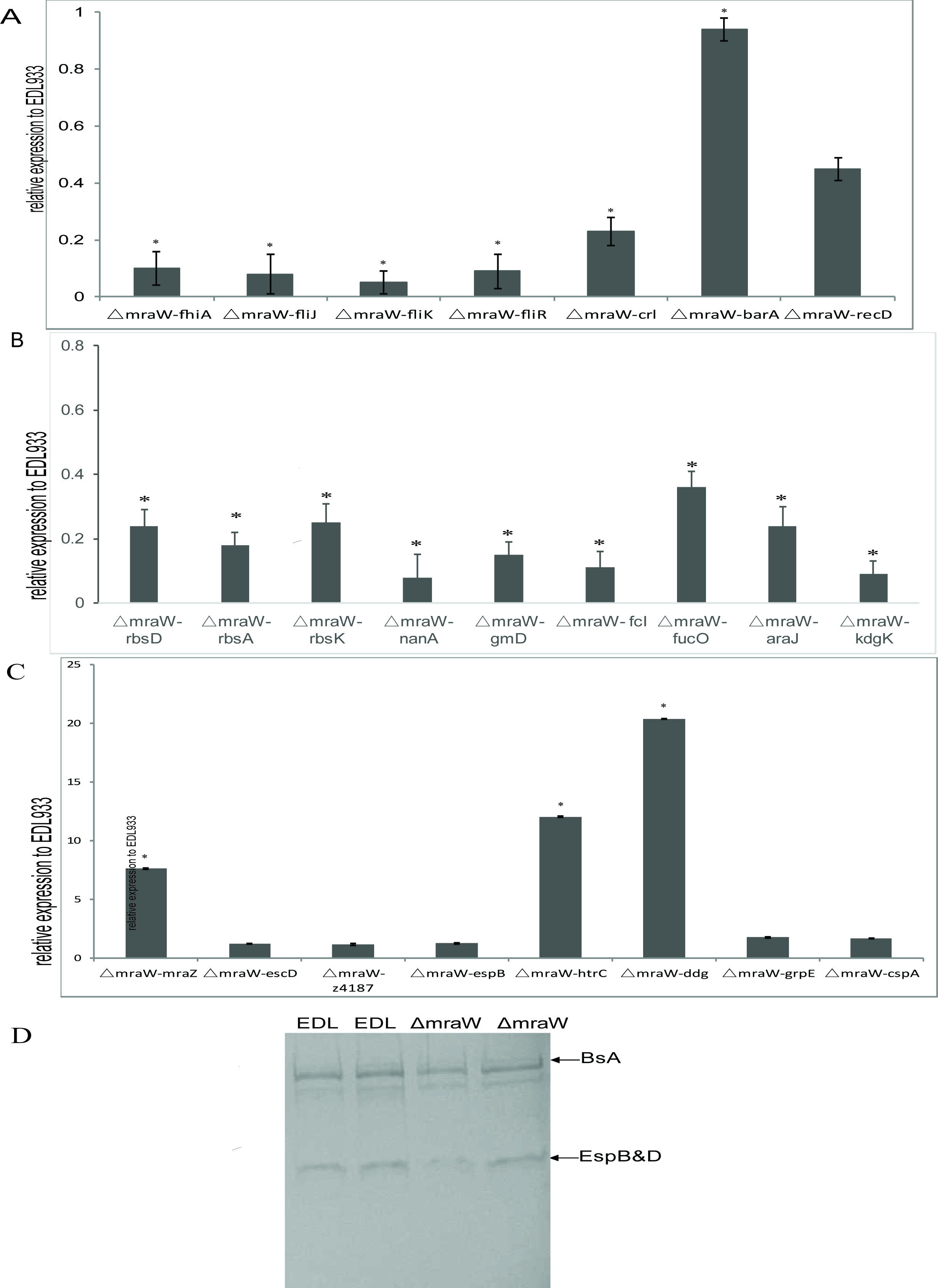
Effect of *mraW* on gene expression at transcriptional level and on T3S profile. Relative mRNA expression of selected genes was normalized to that of the housekeeping gene *gapA*. Results represent mean values ± standard deviations (SD) for three independent experiments. Differences were analyzed for significance using T-test with significant difference between two strains (P <0.01). (A) Motility genes including *fhiA, fliJ, fliK, fliR, crl, barA* and *recD* that were downregulated in mRNA expression compared to wild type strain EDL933. (B) Metabolic genes including *rbsD, rbsA, rbsK, nanA, gmD, fcI, agaI-2, galP, fucO, araJ*, and *kdgK* that were upregulated in mRNA expression compared to wild type strain EDL933. The expression of the gene in EDL933 was set to 1. (C) Genes including *mraZ, escD, z4187, espB, htrC, ddg, grpE* and cspA that were upregulated in mRNA expression compared to wild type strain EDL933. The expression of the gene in EDL933 was set to 1. (D) SDS-PAGE gels showing T3 Secretion profile for EDL933 and mraW deletion mutant strain. BSA and EspB/D bands were indicated. BSA was loaded as control to rule out the precipitation process as a source of variation and to act as a co-precipitant. Strains were cultured in MEM-HEPES to an OD600 of 0.8 and culture supernatants were TCA-precipitated, separated by SDS-PAGE and stained with Colloidal blue.

Due to the antagonistic functions of *mraW* and *mraZ* and their effect on virulence, the expression of *mraZ* was also examined. *mraZ* expression showed more than 7 times increase in the *mraW* mutant compared to the wild type, confirming the antagonistic effect previously observed (19) (Fig. 5D).

## DISCUSSSION

In this study, we investigated the role of MraW in genome-wide DNA methylation and its role in virulence in *E. coli* O157:H7. We found the *mraW* deletion mutant has a genome wide effect on methylation levels of a large number of genes including flagellar related genes, metabolic and respiratory pathway genes, curli related genes, T3SS genes and other virulence related genes. We also found that reduced methylation was correlated altered expression of 21 of the 24 genes tested. MraW was found to have a profound effect on the transcription and protein expression of flagellar genes and motility. MraW was also found to be able to bind to DMRs of flagellar genes and their promoters and methylate cytosine *in vitro*.

The rRNA MTases have been mainly involved in ribosome biogenesis and translation fidelity (20, 21). Some rRNA MTases have been reported to play a role in antibiotic resistance and stress response (22–30). The 16S rRNAMTase encoded by *mraW* is known to be involved in cell division and PG synthesis (12). Recently, *mraW* (*rsmH*) was shown to increase the virulence of *S. aureus in vivo* (13). It was found that the methylation of C1412 of the 16S rRNA which corresponds to C1402 of 16S rRNA in *E. coli* enhanced the animal lethality of *S. aureus* due to increased resistance to oxidative stress in host animals (13). These observations suggest that *mraW* has a wider role than 16S rRNA methylation and cell division.

MraW participated in DNA methylation at whole genome level. A total of 219 promoters and 336 genes showed a decreased methylation level in Δ*mraW*. The DNMT activity of MraW was detected *in vitro* using a cytosine methylation assay, confirming its DNA methyltransferase function. A linear correlation between DNMT activity and protein concentration was found. In addition, we found that MraW can directly bind DNA sequences with DMRs, suggesting that MraW can directly bind to DNA. The default substrate of MraW is the 30S subunit when it functions as an RNA MTase. The crystal structure of MraW shows two distinct but structurally related domains: the typical MTase domain and the putative substrate recognition and binding domain (31). It remains to be investigated whether the substrate recognition and binding domain recognises DNA. The MraW crystal structure contains a cytidine binding site (31). Eight hydrophobic residues including Trp139, Ala143, Ala146, Tyr155, Trp211, Val212, Ile148 and Leu152 interacts indirectly with cytidine forming a hydrophobic environment via a water molecule. Therefore, there is a possibility that the cytosine from DNA sequences could be recognized and methylated by MraW.

Our results showed that MraW affected flagellar motility directly through regulation of the expression of flagellar genes. A potential binding motif and direct binding were found between MraW protein and flagellar related nucleotide sequences including *fliJ, fliR, fhiA*, and *fliK*, indicating a direct physical interaction between MraW and these DNA sequences. The flagellum is composed of three parts: the basal body, the hook, and the filament (32, 33). FliR is one of the six integral membrane proteins of the export apparatus. FliJ participates in forming an ATPase ring complex at the export gate that also plays an important role in substrate recognition with FliI (34, 35). FliK takes part in switching flagellar secretion mode from hook construction to filament elongation and controlling the length of the hook (36). The diminished flagella on the cell surface in the *mraW* mutant is possibly related to the repression of *fliK*. *fhiA* is a homolog of *flhA* which coordinates the export of flagellins (37, 38). The methylation level of *qseB* was also decreased in the mutant. However, QseB represses the motility only in the absence of QseC. Taken together, *mraW* exerts its effect on flagellin production/secretion and motility by acting on the basal apparatus, hook construction, filament length and secreted proteins.

From the mouse colonization results, we found a decreased level of colonization by the *mraW* mutant both using BALB/c mice, a model firstly developed by Mohawk *et al* (18, 39). A lower colonization by the *mraW* mutant is most likely due to nutritional constraints. Carbon nutrition metabolism was found to have essential effect in the mouse colonization of *E. coli* (10, 40) and sialic acid, ribose, mannose and fucose are involved in the colonization of *E*. *coli* in mice (10, 11). We found many genes that were decreased both in methylation and transcriptional levels in the *mraW* mutant are related to energy metabolism mainly related to carbon nutrition metabolism including *rbsD, rbsA*, and *rbsK* for ribose metabolism, *nanA* for sialic acid, *gmD* for mannose, *fcI* and *fucO* for fucose. Other carbon metabolism genes include *agaI-2* for isomerase, *araJ* for arabinose, and *kdgK* for hexuronates. In addition, the methylation and levels of *crl* were reduced in the *mraW* mutant compared to the wild type. As an activator of curli production, Crl binds to stationary phase sigma subunit of RNA polymerase directly (41). Hence a reduced expression of *crl* might affect colonization as well via curli which can assist the colonization of *E*. *coli* (42). The decreased colonization by the *mraW* mutant is less likely due to reduction of motility as less or non-motile *E. coli* O157:H7 was preferred in the animal intestine including mice (11, 43).

Overall, we found a positive correlation between alteration of methylation levels in genes or promotors and levels of transcription except T3SS genes in the *mraW* mutant, suggesting that *mraW* plays a role in genome wide DNA methylation either directly or indirectly and consequently affects transcription and function of a large number of genes. There are a number of possible mechanisms. Firstly, MraW has a dual function as a DNA methylase directly methylate DNA in regions where it can bind. A direct binding of MraW protein and flagellar coding or promoter regions were found indicating a physical interaction between MraW and DNA sequences. Common motifs were also found in subset of genes such as a motif in the 6 DMRs from 3 genes *fliJ, fliK* and *fhiA.* However no common motifs were found across the large number of genes that have been affected. Secondly MraW may have affected the expression of other methylases that in turn affected the methylation of other genes. Thirdly, the effect of MraW might be achieved via MraZ. The deletion of *mraW* perturbs *mraZ* which is antagonistic to *mraW* and a known regulator of *mraW* and 100 other genes (61 activated and 31 repressed) in *E. coli* K-12 (19). However, when *mraZ* was mildly over produced it affected the expression of 970 genes in *E. coli* K-12 (19). Hence, *mraW* might control genome wide gene expression via the antagonistic relationship with *mraZ*. We found the expression of *mraZ* was increased in the *mraW* mutant which is consistent with the observations in *E. coli* K-12 (19).

Although the vast majority of the differentially methylated genes/promoters had reduced methylation in the *mraW* mutnat, 13 showed increased methylation levels. A plausible explanation for these small number of genes with increased methylation is that these DMRs may be methylated by other methylases that were repressed by *mraW*. It is unlikely all of these were false positives as two were confirmed by BSP PCR sequencing. Further studies will be required to elucidate the mechanisms involved.

In conclusion, we found that *mraW* plays a role in gene regulation through DNA methylation in addition to known function in methylating 16S rRNA to increase mRNA decoding fidelity. *mraW* affected DNA methylation, motility, and mouse colonization and clearly plays a role in virulence in *E. coli*.

## MARERIALS AND METHODS

### Bacterial Strains, Plasmids and Bacterial culture

Details of bacterial strains and plasmids used in this study are listed in Table S4. Bacteria were routinely cultured in Luria-Bertani (LB, Miller) broth or agar. Antibiotics were included when required at the following concentrations: 100 µg ml^−1^ ampicillin, 50 µg ml^−1^ kanamycin and 50 µg ml^−1^ chloramphenicol. Other chemicals added to media were 0.2% L-arabinose, bromo-chloro-indolyl-galactopyranoside (X-gal) 20 μg ml^−1^ (dissolved in dimethylformamide) and Isopropyl β-D-1-thiogalactopyranoside (IPTG) 2μg ml^−1^. Most of motility related experiments were performed at an OD of 0.6 at which density bacterial growth is in the log phase.

### Construction of *mraW* deletion mutant in EDL933

Construction of the *mraW* deletion mutant in EDL933 was performed using one step method as described by Datsenko and Wanner (44). The *kmr* gene was amplified by PCR from plasmid pRS551 (45) with primer pair P1 and P2 (Table S5) (46). EDL933ΔmraW mutant was confirmed by PCR and sequencing. The primer pairs P3 and P4, P5 and P6 (Table S5) were used to confirm *mraW* gene deletion.

### Construction of plasmids for expression and complementation

Complementation plasmid pBADmraW was constructed using PCR product amplified with high fidelity Phusion polymerase cloned into pBAD/Myc-HisA (Table S4). Primers including 5-mraWF and 3-mraWR were used (Table S5). *E. coli* strain DH5a was used as the intermediate host strain for cloning and all constructs were verified by sequencing.

### Motility assays

Swimming motility was evaluated as described by Xu and Xu (7). Briefly, wild type, the *mraW* mutant and complement strains were cultured overnight and stab inoculated with a sterile inoculating needle and incubated at 37°C for 16h. All strains were tested in triplicate and each experiment was carried out on three separate occasions. The motility radius of each strain was measured and analyzed by t-Test.

### Transmission electron microscopy

TEM was carried out as previously described (7). In brief, EDL933 and its mutant EDL933ΔmraW were cultured in LB broth with shaking till optical density (OD600) of 1.0. Bacteria on TEM grids were stained with 1% (wt/vol) phosphotungstic acid and examined with a Philips Tecnai 12 transmission electron microscope. Images were obtained with Gatan Digital Micrograph Imaging System by an Erlangshen CCD camera (Gantan).

### Preparation of H7 supernatant protein

To examine the H7 Flic protein expression in the wild type and the mutant strains, bacteria were cultured overnight in LB at 37°C with shaking and sub-cultured in a dilution of 1:100 in LB. Strains were grown to a final OD600 of 1.0 and then centrifuged at 4000g for 30min at 4°C and supernatants were filtered through 0.45 mm low protein-binding filters (Millipore). A 10% (v/v) final concentration of trichloroacetic acid (TCA; Sigma-Aldrich) was used to precipitate the proteins. Supernatants were incubated overnight at 4°C for thorough precipitation and centrifuged at 4000g for 30min at 4°C. Protein pellets were air-dried and resuspended in an appropriate volume of resuspension buffer (1.5M Tris-HCL) in order to standardize samples and to take into account the slight variation in OD600 at which cultures were harvested.

### H7 Flic protein detection

Supernatant protein was resolved on a 12% sodium dodecyl sulfate-polyacrylamide (SDS-PAGE) gel. SDS-PAGE separated proteins were transferred onto Hybond ECL nitrocellulose membrane (Amersham Biosciences) with a Trans-Blot electrophoretic transfer cell (Bio-Rad). Nitrocellulose membranes were blocked with 8% (w/v) dried milk powder (Marvel) in PBS at 4°C overnight and incubated with the relevant antibodies diluted in wash buffer 0.05% (v/v) polyoxyethylene sorbitan monolaurate (Tween 20, Sigma-Aldrich) in PBS at the following dilutions: primary antibodies against H7 flagellin (Statens Serum Institut, Denmark) were diluted 1:1000 and secondary antibody with IR Dye 800-labeled anti-rabbit IgG (Rockland, Gilbertsville, PA, U.S.) were diluted 1:10000. Fluorescence signal was captured by Odyssey imager (LI-COR).

### DNA sequencing, library preparation and genome methylation sequencing

Genomic DNA was extracted with Wizard Genomic^®^ DNA Purification Kit (Promega). DNA was sonicated to 100-300 bp fragments. Bisulfate treatment was carried out by ZYMO EZ DNA Methylation-Gold kit (Zymo Research). By gel purification, DNA fragments with proper size were used for PCR amplification. The proper size PCR-amplified fragments were sequenced using HiSeq2000. Sequencing data was mapped onto the reference *E. coli* O157:H7 strain EDL933 to obtain the genome methylation data.

### Validation of genome bisulfite sequencing results by bisulfite sequencing PCR

Genomic DNA bisulfate treatment was carried out by EpiTect^®^ Bisulfite kit (Qiagen). Primers for Bisulfite sequencing PCR were designed using Methyl Primer Express v1.0 (Table S5, from fhiAP1F to treR2R). PCR was carried out using the DNA template which was treated by EpiTect^®^ Bisulfite kit. PCR product was cloned into pMD-18-T and sequenced. For each DNA fragment, at least 10 clones were selected for sequencing to minimize the sequencing error. Cytosine methylation data was achieved by mapping the sequencing data to the reference *E. coli* O157:H7 str. EDL933.

### RNA extraction, cDNA synthesis and quantitative reverse transcription PCR

Overnight cultures of *E. coli* EDL933 and, its mutant strain was diluted 100-fold in LB broth and then grown to an OD600 of 0.6 with shaking. Total RNA was extracted using RNeasy Mini Kit (Qiagen) following the manufacturer’s instructions. RNA was treated with DNase I (NEB). Expression of *fhiA, fliJ, fliK, fliR, crl, barA, recD, mraZ, escD, z4187, espB, htrC, ddg, gprE, cspA rbsD, rbsA, rbsK, nanA, gmD, fcI, agaI-2, galP, fucO, araJ*, and *kdgK* were quantified by quantitative reverse transcription-PCR (qRT-PCR) analysis. Reverse transcription was performed using PrimeScript^®^ RT reagent Kit (Perfect Real Time) (TaKaRa). qRT-PCR was carried out using SYBR^®^ Premix Ex TaqTM II (Perfect Real Time) (TaKaRa) using a Rotor-Gene Q thermal cycler (QIAGEN). Data was analyzed with Rotor-Gene Q Series Software, version 1.7 (QIAGEN). Data were normalized to the endogenous reference gene *gapA* and analyzed by the cycle threshold method (2-ΔΔCT) (47). Three independently isolated cDNA samples were analyzed. Primers for amplifying *gapA, fhiA, fliJ, fliK, fliR,crl, barA, recD, mraZ, escD, z4187,espB, htrC, ddg, gprE, cspA rbsD, rbsA, rbsK, nanA, gmD, fcI, agaI-2, galP, fucO, araJ*, and *kdgK* are detailed in Table S5 (From fhiA RT F to kdgK RT R).

### EMSA test

The promoter of *fhiA* & *fliK*, and genes of *fliJ* & *fliR* were synthesized and cloned into pUC19-c vector with DH10b competent cell. As the sequence of *fliR* gene exceeds 500 bp, it was divided two frarments. The cloned sequences and constructed plasmids were listed in Table S4. Plasmid pDS17039, pDS17040, pDS17041, pDS17042 and pDS17043 were achieved respectively from the sequence of *fhiA, fliK, fliJ* and *fliR*. For preparation of fluorescent FAM labeled probes, the promoter region of pDS17039 plasmid, pDS17040 plasmid, pDS17041 plasmid, pDS17042 plasmid and pDS17043 plasmid, was PCR amplified with Dpx DNA polymerase using primers of M13F (FAM) and M13R. The FAM-labeled probes were purified by the Wizard^®^ SV Gel and PCR Clean-Up System (Promega, USA) and were quantified with NanoDrop 2000C (Thermo, USA). MraW protein were prepared and purified as previously described (Zhang Heng). EMSA was performed in a 20 µl reaction volume that contains 40 ng probe and varied of MR proteins, in a reaction buffer of 50 mM Tris-HCl [pH 8.0], 100 mM KCl, 2.5 mM MgCl2, 0.2 mM DTT, 2 μg salmon sperm DNA and 10% glycerol. After incubation for 20 min at 25 °C, the reaction system was loaded into 2% TBE gel buffered with 0.5×TBE. Gels were scanned with Image Quant LAS 4000 mini (GE Healthcare).

### DNMT activity test

MraW protein were prepared and purified as previously described (Zhang Heng). DNMT activity test was performed in accordance with EpiQuik(™) DNMT Activity/Inhibition Assay Ultra Kit manual (Colorimetric) (Epigentek). DNMT activity was calculated as the following formula:

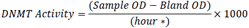

* Incubation time of protein and DNMT substrate (cytosine).

### Identification of MraW key motifs

To examine the potential interacting motif of *mraW* in whole genome, flagellar and heat shock genes, the motifs in all DMRs were searched by meme 4.8.1 software. Motifs with p value < 0.0001 were selected.

### Mouse infection studies

All animal work was approved by Laboratory Animal Welfare & Ethics Committee at Chinese National Institute for Communicable Disease Control and Prevention. To construct selective bacteria maintaining *in vivo* in the absence of antibiotic pressure, a constitutive plasmid pGEN-luxCDABE encoding bacterial luciferase was electro-transformed into the wild type and the *mraW* deletion mutant (48). Conventional mice model was successfully used for intestinal colonization with high doses of *E*. *coli* O157:H7 (39). Six-week-old female BALB/c mice from Charles River (China, Beijing) was infected by gavage with 10^9^ and 10^10^ CFU of the wild type and the mutant both containing pGEN-luxCDABE. To prepare the inoculum, overnight culture at 37 °C with shaking at 225 rpm in LB media with 100 µg/ml ampicillin were pelleted by centrifugation for 10 min at 1,000 × g. The bacteria were then resuspended in 1/40th volume of PBS. Groups of 10 mice each were gavaged with one hundred microliters of the organisms. The day of infection was designated as day 0. The shedding extent of individual mice was monitored as below. Fecal pellets from each mouse were collected, weighed, and suspended 1:10 (wt/vol) in PBS. The fecal pellet and PBS mixtures were mixed by vertexing at room temperature for 1 min, every 10 min, for 30 min. Ten-fold dilutions of the mixtures were made with PBS, and two aliquots each containing one hundred microliters of the homogenates were plated onto Sorbitol–MacConkey agar plates containing 100 µg ml^−1^ ampicillin and 50 µg ml^−1^ kanamycin respectively. Plates were incubated overnight at 37 °C. Colonies with luminescence were imaged by using In-Vivo FX Pro (Bruker) and counted the next day.

**Table 1.**
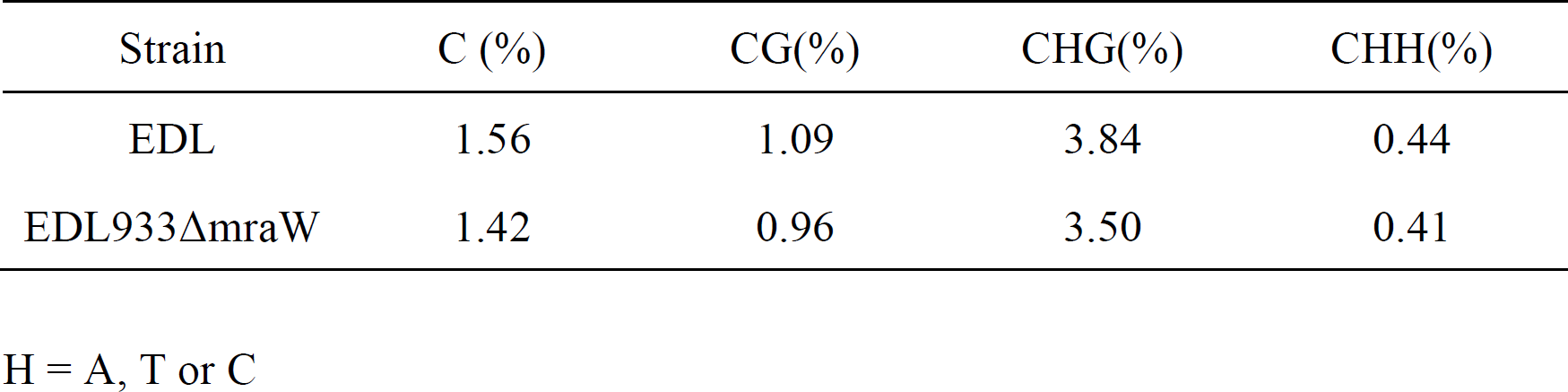
Whole genome methylation level of C, CG, CHG and CHH

## SUPPLEMENTAL MATERIAL

TABLE S1

TABLE S2

TABLE S3

TABLE S4

TABLE S5

## Acknowledgements

This work was supported by grants from the National Natural Science Foundation of China (81471918, 31370166 and 81772152) and the State Key Laboratory for Infectious Disease Prevention and Control (2014SKLID204).

## References

1. Garg AX, Clark WF, Salvadori M, Macnab J, Suri RS, Haynes RB, Matsell D, Walkerton Health Study I. 2005. Microalbuminuria three years after recovery from *Escherichia coli* O157 hemolytic uremic syndrome due to municipal water contamination. Kidney Int 67:1476–1482.

2. Friedrich AW, Zhang W, Bielaszewska M, Mellmann A, Kock R, Fruth A, Tschape H, Karch H. 2007. Prevalence, virulence profiles, and clinical significance of Shiga toxin-negative variants of enterohemorrhagic *Escherichia coli* O157 infection in humans. Clin Infect Dis 45:39–45.

3. Xu X, McAteer SP, Tree JJ, Shaw DJ, Wolfson EB, Beatson SA, Roe AJ, Allison LJ, Chase-Topping ME, Mahajan A, Tozzoli R, Woolhouse ME, Morabito S, Gally DL. 2012. Lysogeny with Shiga toxin 2-encoding bacteriophages represses type III secretion in enterohemorrhagic *Escherichia coli*. PLoS Pathog 8:e1002672.

4. Friedland JA, Herman TE, Siegel MJ. 1995. *Escherichia coli* O157:H7-associated hemolytic-uremic syndrome: value of colonic color Doppler sonography. Pediatr Radiol 25 Suppl 1:S65–67.

5. Dobbin HS, Hovde CJ, Williams CJ, Minnich SA. 2006. The *Escherichia coli* O157 flagellar regulatory gene flhC and not the flagellin gene fliC impacts colonization of cattle. Infect Immun 74:2894–2905.

6. Bretschneider G, Berberov EM, Moxley RA. 2007. Reduced intestinal colonization of adult beef cattle by *Escherichia coli* O157:H7 tir deletion and nalidixic-acid-resistant mutants lacking flagellar expression. Vet Microbiol 125:381–386.

7. Xu Y, Xu X, Lan R, Xiong Y, Ye C, Ren Z, Liu L, Zhao A, Wu LF, Xu J. 2013. An O island 172 encoded RNA helicase regulates the motility of *Escherichia coli* O157:H7. PLoS One 8:e64211.

8. Laaberki MH, Janabi N, Oswald E, Repoila F. 2006. Concert of regulators to switch on LEE expression in enterohemorrhagic *Escherichia coli* O157:H7: interplay between Ler, GrlA, HNS and RpoS. Int J Med Microbiol 296:197–210.

9. Chilcott GS, Hughes KT. 2000. Coupling of flagellar gene expression to flagellar assembly in *Salmonella enterica serovar typhimurium* and *Escherichia coli*. Microbiol Mol Biol Rev 64:694–708.

10. Chang DE, Smalley DJ, Tucker DL, Leatham MP, Norris WE, Stevenson SJ, Anderson AB, Grissom JE, Laux DC, Cohen PS, Conway T. 2004. Carbon nutrition of *Escherichia coli* in the mouse intestine. Proc Natl Acad Sci U S A 101:7427–7432.

11. Leatham MP, Stevenson SJ, Gauger EJ, Krogfelt KA, Lins JJ, Haddock TL, Autieri SM, Conway T, Cohen PS. 2005. Mouse intestine selects nonmotile flhDC mutants of *Escherichia coli* MG1655 with increased colonizing ability and better utilization of carbon sources. Infect Immun 73:8039–8049.

12. Kimura S, Suzuki T. 2010. Fine-tuning of the ribosomal decoding center by conserved methyl-modifications in the *Escherichia coli* 16S rRNA. Nucleic Acids Res 38:1341–1352.

13. Kyuma T, Kimura S, Hanada Y, Suzuki T, Sekimizu K, Kaito C. 2015. Ribosomal RNA methyltransferases contribute to *Staphylococcus aureus* virulence. FEBS J 282:2570–2584.

14. Furst AL, Muren NB, Hill MG, Barton JK. 2014. Label-free electrochemical detection of human methyltransferase from tumors. Proc Natl Acad Sci U S A 111:14985–14989.

15. Hughes DT, Clarke MB, Yamamoto K, Rasko DA, Sperandio V. 2009. The QseC adrenergic signaling cascade in Enterohemorrhagic *E. coli* (EHEC). PLoS Pathog 5:e1000553.

16. Pernestig AK, Georgellis D, Romeo T, Suzuki K, Tomenius H, Normark S, Melefors O. 2003. The *Escherichia coli* BarA-UvrY two-component system is needed for efficient switching between glycolytic and gluconeogenic carbon sources. J Bacteriol 185:843–853.

17. Zumbrun SD, Melton-Celsa AR, Smith MA, Gilbreath JJ, Merrell DS, O’Brien AD. 2013. Dietary choice affects Shiga toxin-producing *Escherichia coli* (STEC) O157:H7 colonization and disease. Proc Natl Acad Sci U S A 110:E2126–2133.

18. Mohawk KL, Melton-Celsa AR, Zangari T, Carroll EE, O’Brien AD. 2010. Pathogenesis of *Escherichia coli* O157:H7 strain 86-24 following oral infection of BALB/c mice with an intact commensal flora. Microb Pathog 48:131–142.

19. Eraso JM, Markillie LM, Mitchell HD, Taylor RC, Orr G, Margolin W. 2014. The highly conserved MraZ protein is a transcriptional regulator in *Escherichia coli*. J Bacteriol 196:2053–2066.

20. Burakovsky DE, Prokhorova IV, Sergiev PV, Milon P, Sergeeva OV, Bogdanov AA, Rodnina MV, Dontsova OA. 2012. Impact of methylations of m2G966/m5C967 in 16S rRNA on bacterial fitness and translation initiation. Nucleic Acids Res 40:7885–7895.

21. Das G, Thotala DK, Kapoor S, Karunanithi S, Thakur SS, Singh NS, Varshney U. 2008. Role of 16S ribosomal RNA methylations in translation initiation in *Escherichia coli*. EMBO J 27:840–851.

22. Toh SM, Xiong L, Bae T, Mankin AS. 2008. The methyltransferase YfgB/RlmN is responsible for modification of adenosine 2503 in 23S rRNA. RNA 14:98–106.

23. Bujnicki JM, Rychlewski L. 2001. Sequence analysis and structure prediction of aminoglycoside-resistance 16S rRNA:m7G methyltransferases. Acta Microbiol Pol 50:7–17.

24. Nichols RJ, Sen S, Choo YJ, Beltrao P, Zietek M, Chaba R, Lee S, Kazmierczak KM, Lee KJ, Wong A, Shales M, Lovett S, Winkler ME, Krogan NJ, Typas A, Gross CA. 2011. Phenotypic landscape of a bacterial cell. Cell 144:143–156.

25. Gustafsson C, Persson BC. 1998. Identification of the rrmA gene encoding the 23S rRNA m1G745 methyltransferase in *Escherichia coli* and characterization of an m1G745-deficient mutant. J Bacteriol 180:359–365.

26. Helser TL, Davies JE, Dahlberg JE. 1971. Change in methylation of 16S ribosomal RNA associated with mutation to kasugamycin resistance in *Escherichia coli*. Nat New Biol 233:12–14.

27. Osterman IA, Sergiev PV, Tsvetkov PO, Makarov AA, Bogdanov AA, Dontsova OA. 2011. Methylated 23S rRNA nucleotide m2G1835 of *Escherichia coli* ribosome facilitates subunit association. Biochimie 93:725–729.

28. Lesnyak DV, Sergiev PV, Bogdanov AA, Dontsova OA. 2006. Identification of *Escherichia coli* m2G methyltransferases: I. the ycbY gene encodes a methyltransferase specific for G2445 of the 23 S rRNA. J Mol Biol 364:20–25.

29. Andersen NM, Douthwaite S. 2006. YebU is a m5C methyltransferase specific for 16 S rRNA nucleotide 1407. J Mol Biol 359:777–786.

30. Basturea GN, Dague DR, Deutscher MP, Rudd KE. 2012. YhiQ is RsmJ, the methyltransferase responsible for methylation of G1516 in 16S rRNA of *E*. coli. J Mol Biol 415:16–21.

31. Wei Y, Zhang H, Gao ZQ, Wang WJ, Shtykova EV, Xu JH, Liu QS, Dong YH. 2012. Crystal and solution structures of methyltransferase RsmH provide basis for methylation of C1402 in 16S rRNA. J Struct Biol 179:29–40.

32. Macnab RM. 2003. How bacteria assemble flagella. Annu Rev Microbiol 57:77–100.

33. Barker CS, Meshcheryakova IV, Inoue T, Samatey FA. 2014. Assembling flagella in Salmonella mutant strains producing a type III export apparatus without FliO. J Bacteriol 196:4001–4011.

34. Minamino T. 2014. Protein export through the bacterial flagellar type III export pathway. Biochim Biophys Acta 1843:1642–1648.

35. Sajo R, Liliom K, Muskotal A, Klein A, Zavodszky P, Vonderviszt F, Dobo J. 2014. Soluble components of the flagellar export apparatus, FliI, FliJ, and FliH, do not deliver flagellin, the major filament protein, from the cytosol to the export gate. Biochim Biophys Acta 1843:2414–2423.

36. Uchida K, Aizawa S. 2014. The flagellar soluble protein FliK determines the minimal length of the hook in *Salmonella enterica serovar Typhimurium*. J Bacteriol 196:1753–1758.

37. Bange G, Kummerer N, Engel C, Bozkurt G, Wild K, Sinning I. 2010. FlhA provides the adaptor for coordinated delivery of late flagella building blocks to the type III secretion system. Proc Natl Acad Sci U S A 107:11295–11300.

38. Ibuki T, Uchida Y, Hironaka Y, Namba K, Imada K, Minamino T. 2013. Interaction between FliJ and FlhA, components of the bacterial flagellar type III export apparatus. J Bacteriol 195:466–473.

39. Mohawk KL, O’Brien AD. 2011. Mouse models of *Escherichia coli* O157:H7 infection and shiga toxin injection. J Biomed Biotechnol 2011:258185.

40. Peekhaus N, Conway T. 1998. What’s for dinner?: Entner-Doudoroff metabolism in *Escherichia coli*. J Bacteriol 180:3495–3502.

41. Bougdour A, Lelong C, Geiselmann J. 2004. Crl, a low temperature-induced protein in *Escherichia coli* that binds directly to the stationary phase sigma subunit of RNA polymerase. J Biol Chem 279:19540–19550.

42. Lloyd SJ, Ritchie JM, Rojas-Lopez M, Blumentritt CA, Popov VL, Greenwich JL, Waldor MK, Torres AG. 2012. A double, long polar fimbria mutant of *Escherichia coli* O157:H7 expresses Curli and exhibits reduced in vivo colonization. Infect Immun 80:914–920.

43. Sharma VK, Casey TA. 2014. *Escherichia coli* O157:H7 lacking the qseBC-encoded quorum-sensing system outcompetes the parental strain in colonization of cattle intestines. Appl Environ Microbiol 80:1882–1892.

44. Datsenko KA, Wanner BL. 2000. One-step inactivation of chromosomal genes in *Escherichia coli* K-12 using PCR products. Proc Natl Acad Sci U S A 97:6640–6645.

45. Simons RW, Houman F, Kleckner N. 1987. Improved single and multicopy lac-based cloning vectors for protein and operon fusions. Gene 53:85–96.

46. Oka A, Sugisaki H, Takanami M. 1981. Nucleotide sequence of the kanamycin resistance transposon Tn903. J Mol Biol 147:217–226.

47. Livak KJ, Schmittgen TD. 2001. Analysis of relative gene expression data using real-time quantitative PCR and the 2(-Delta Delta C(T)) Method. Methods 25:402–408.

48. Lane MC, Alteri CJ, Smith SN, Mobley HL. 2007. Expression of flagella is coincident with uropathogenic *Escherichia coli* ascension to the upper urinary tract. Proc Natl Acad Sci U S A 104:16669–16674.

